# Accessible and accurate cytometry analysis using fluorescence microscopes

**DOI:** 10.1101/2025.01.22.634380

**Authors:** Daniel Foyt, Yiming Kuang, Samma Rehem, Klaus Yserentant, Bo Huang

**Affiliations:** UCSF-UC Berkeley Joint Graduate Program in Bioengineering, University of California San Francisco, San Francisco, California, 94143, United States of America; Department of Pharmaceutical Chemistry, University of California San Francisco, San Francisco, California, 94143, United States of America; Barnard College, New York City, New York, 10027, United States of America; Department of Biochemistry and Biophysics, University of California San Francisco, San Francisco, California, 94143, United States of America; Chan Zuckerberg Biohub San Francisco, San Francisco, California, 94158, United States of America

## Abstract

We have developed a method along with a python-based analysis tool to capture images and produce flow cytometry like data utilizing simple accessible microscopes. Utilizing the recently developed generalist algorithms for cell segmentation, our approach easily segments semi-adherent or suspended cells facilitating quantification of fluorescent intensity similar to flow cytometry. We have shown that our approach exhibits similar speed and enhanced sensitivity when compared to typical flow cytometry. The utility of our approach is demonstrated by screening a set of 88 prime editing conditions utilizing the integration of mNeonGreen_11_ as a reporter.

## Introduction

Measurements of the number, size, physical phenotype, and presence or absence of a specific protein or reporter in a pool of cells are a crucial part of a biologist tool kit. Currently, these functions are typically carried out using flow cytometry. Flow cytometry operates by flowing single cells in a steady aligned stream using fluid flow techniques and analyzing droplets or cells as they pass through a laser or electrical field (1). Flow cytometry has been a very powerful method in the field of immunology and molecular biology to measure the fluorescent readout of thousands of cells (1). While powerful in its throughput, the information obtained from this technique is inherently lacking in depth as it typically only provides an average intensity of the fluorescent reporter while not providing spatial information about the signal distribution in the context of other cellular components. Additionally, the necessity to suspend cells in solution for analysis can be limiting in some cases. Historically, more detailed measurements of cellular physical phenotypes were made with microscopy, which would necessitate compromising the scale and speed of the assay compared to flow cytometry. Moreover, segmentation of images so that individual cells growing in groups can be distinguished is challenging without explicit reporters for cell boundaries(2,3). With the advancement of microscope automation, image analysis tools, and the availability of suitable microscopes to many biologists, microscopy-based cytometry could be come a more accessible alternative to flow cytometry.

## Results

### Development and validation of microscopy-based cytometry

One of the main impediments to a microscopy-based cytometry method for adherent cells is the ability to segment individual cells reliably when they grow in a cluster. While there are algorithms developed for this task, such as Cellpose(4), a generalist deep learning-based segmentation method, they do not always reliably define cell boundaries in bright-field images due to cells physically overlapping and lack of contrast for cell-cell interfaces(2,3). Although fluorescent staining for cell boundaries (such as plasma membrane stain) can facilitate segmentation, it consumes a color channel of the microscope. Therefore, we sought to develop an approach for reliable segmentation using bright-field images only. We reasoned that inducing cells to partially detach from one another and the surface would reduce their overlap, making the images easier to segment. To determine if partially detached cells facilitate better segmentation, we first imaged cells that were adhered to the surface of a coverglass-bottom 96- well plate using a Nikon Ti-E inverted wide-field microscope and attempted to segment the images using Cellpose(4). As expected, brightfield segmentations match poorly with fluorescence images from nuclear staining, which served as the ground truth control (Fig. 1a). We then treated the cells in the well with trypsin, which is used in flow-cytometry analysis of adherent cells to detatch the cells. We observed that, with a brief trypsin treatment, cells stayed attached to the surface but that the boundaries between cells were much more apparent, improving the segmentation results (Fig. 1b). To see if the complete detachment of cells from the surface would further improve the discrimination of cell boundaries, we treated cells with trypsin again, mechanically agitated the media to detach the cells (similar to the generation of cell suspension for common flow cytometry preparation procedure), and then centrifuged the well plate to let the cells settle down to the surface for image capture. Segmentation of these images resulted in only a slight improvement in cell segmentation (Fig. 1c). For quantitative assessment, we compared the number of segments obtained from the brightfield images with the number of nuclei counted. Treating cells with trypsin alone and trypsin plus resuspension substantially improved the accuracy compared to adherent conditions (Fig. 1d). Additionally, these treatments decreased the false negative (Fig. 1e) and false positive (Fig. 1f) ratios for cell detection.

**Fig. 1.**
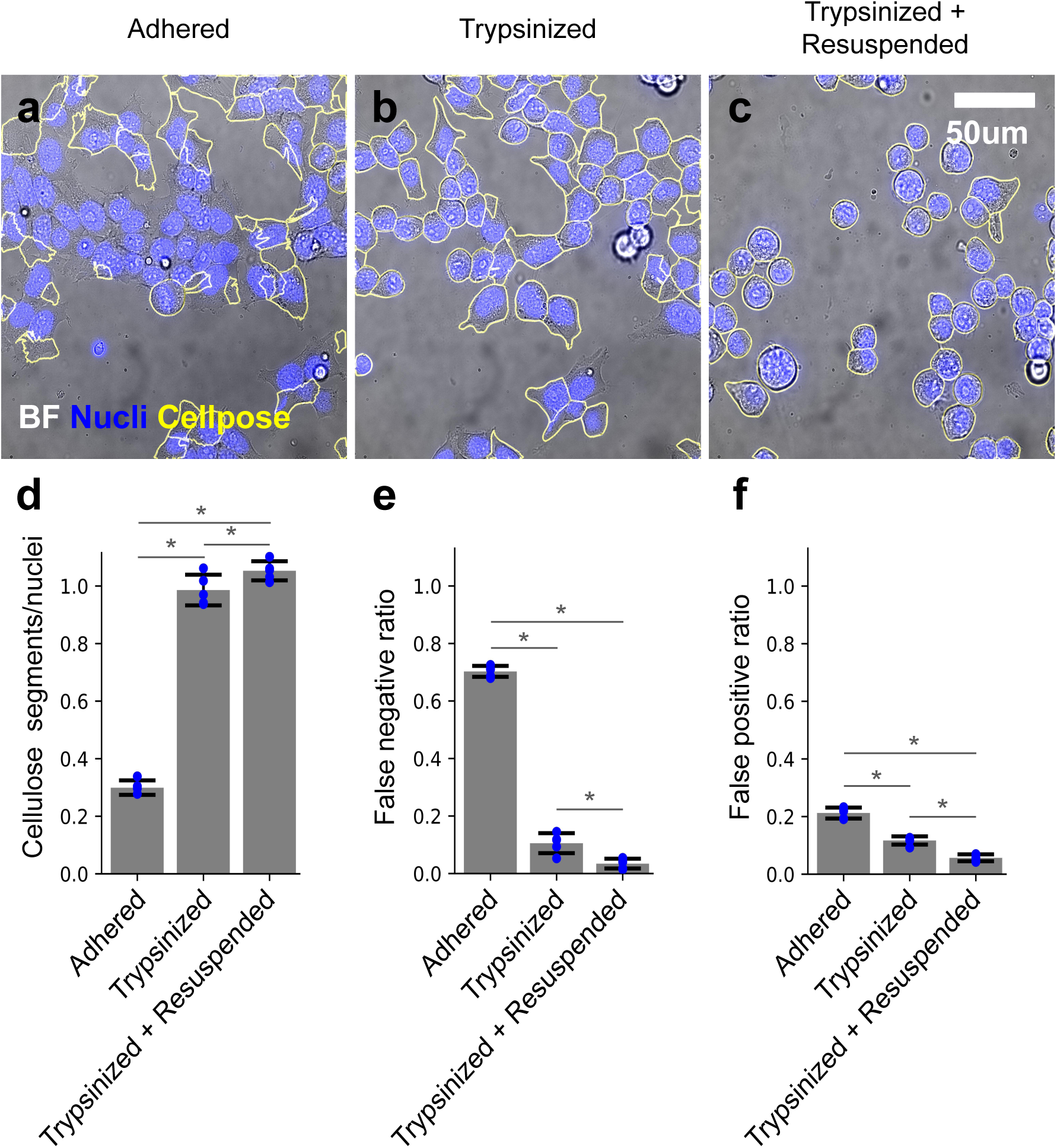
Trypsin treatment improves Cellpose segmentation of brightfield images for adherent cells. **a**. A representative brightfield image of adherent HEK cells with Cellpose segment outlines highlighted, overlaid with a fluorescence image of the nuclear stain in blue color. **b**. Same as **a** for trypsinized cells. **c**. Same as **b** for trypsinized and resuspended cells. **d**. Cellpose segments per number of counted nuclei. **e**. False negative ratio of Cellpose. **f**. False positive ratio of Cellpose. Error bars represent standard deviation across 4 independent experiments. * denotes p < 0.05 when comparing groups.

Cellpose utilizes two parameters when determining the boundary of a cell; the first is an estimated segment diameter, as the models used were trained on scaled images with constant diameters, which may be different from the experimental data. The second is a flow threshold, which is the maximum allowed error of the flows determined by the model and the predicted segments. To ensure we utilized the optimal parameters in Cellpose and that the difference in segmentation accuracy was not due to improper parameter selection, we screened a combination of 25 diameters ranging from 25 to 505 pixels and 21 flow threshold values from 0.05 to 2.05 for a total of 525 parameter combinations. We calculated the number of segments obtained from the Cellpose segmentation of brightfield images as a fraction of the number of nuclei counted. We observed that for untreated adherent cells, Cellpose only produced 30% of the number of segments as there were nuclei with optimal parameters (cell diameter = 145, flow threshold = 0.95-2.05) (Fig. 2a). The Cellpose segments/nuclei measurements for the trypsinized-only cells had a maximum value of close to 1 with optimal parameters of cell diameter = 145 and flow threshold = 0.95-2.05 (Fig. 2b). The optimal parameters (Cellpose segments/nuclei = 1) for trypsinized and resuspended cells were in a slightly wider range. Still, there were combinations of Cellpose parameters (45 < diameter (pixels) < 265, flow threshold > 0.45) that yielded more segments than counted nuclei (Fig. 2c). We also calculated false negative and false positive rates for all parameter combinations and trypsinization conditions. The adhered cell condition had a much higher false negative and false positive ratio compared to the trypsinized and trypsinized and resuspended conditions (Fig. 2d-i).

**Fig. 2.**
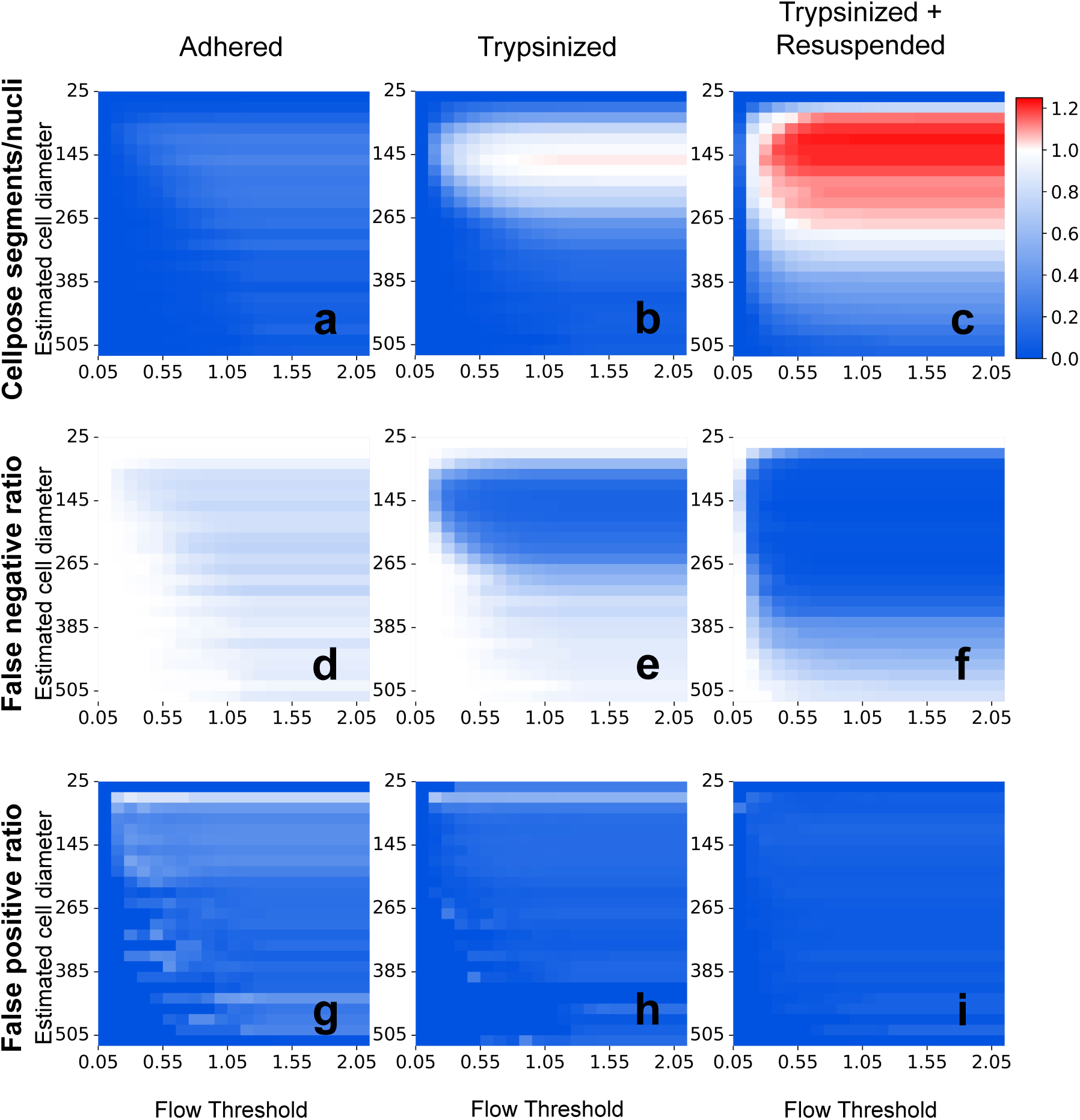
Cellpose parameter optimization. **a-c**. Heat map of Cellpose segments per number of counted nuclei for all combinations of estimated cell diameter and flow threshold assessed for adhered (left), trypsinized (middle), and trypsinized + resuspended (right) condition. **d-f**. Same as **a-c** for false negative ratio. **g-i**. Same as **a-c** for false positive ratio.

To understand the upper limit of the number of cells we could reliably segment and assay in our implementation, we varied the number of cells loaded into a well in a 96-well plate from 15,600 to 500,000, took brightfield images, and segmented the images. Cell densities from 15,600 to 62,500 cells/well were reliably segmented (Fig. 3a), and the number of segments linearly correlated with the number of cells loaded. At higher densities, the segmentation efficiency worsens (Fig. 3b). At densities greater than 62,500 cells/well, the cells no longer form a single layer, and all of the cells can not be reliably segmented (Fig. 3a 125k).

**Fig. 3.**
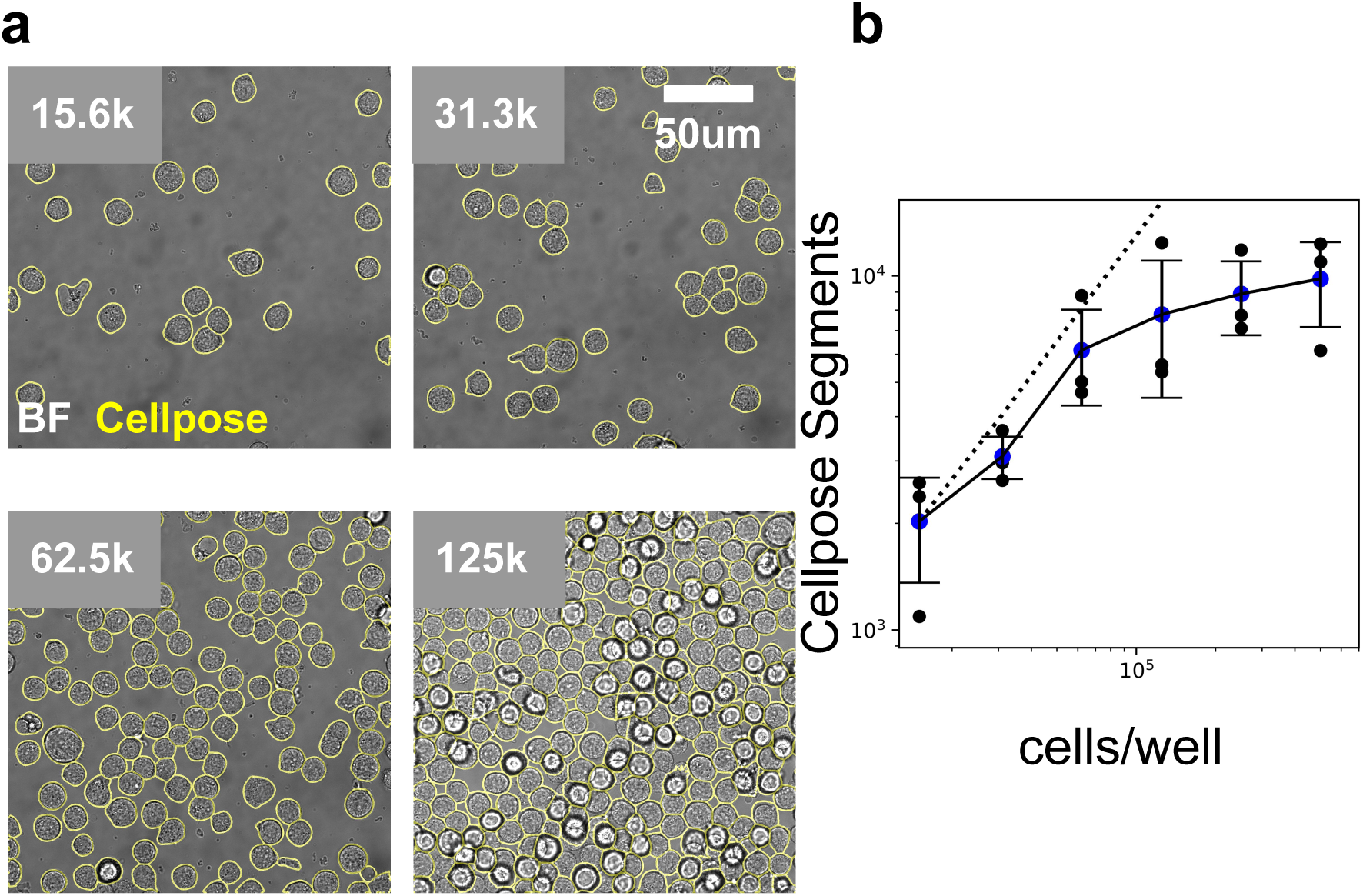
Cell density optimization for Cellpose segmentation. **a**. Representative images and Cellpose segment outlines for 15.6k, 31.3k, 62.5k, and 125k cells/well of a 96 well plate. **b**. Cellpose segments obtained from 49 images for varying number of loaded cells. Dotted line is the theoretical trend expected if Cellpose segments and number of loaded cells were perfectly correlated.

We performed a cell mixing experiment to determine the single-cell purity of the data obtained from our method. We obtained two cell lines, each expressing spectrally district fluorescent proteins; one HEK293 cell line stably expressing mApple created via landing pad integration (5), and a HeLa cell line with HspA8 endogenously tagged with the mNeonGreen2_11_ fragment (6) (mNG2_11_) and stably expressing mNG3A_1-10_ (7) which yields green fluorescent HspA8 protein. We mixed these cell lines in equal proportions, seeded them into wells, and allowed the cells to adhere and grow. We gently trypsinized these cells and captured images in the brightfield, mNG, and mApple channels (Fig. 4a). We observed that 0.90% of cells contained a mix of signals from both mNG and mApple, both above the wild-type control signal, suggesting a doublet rate of 1.80% (from a 50/50 mix of cells, half of the doublets will arise from two cells with the same fluorescence protein) (Fig. 4b).

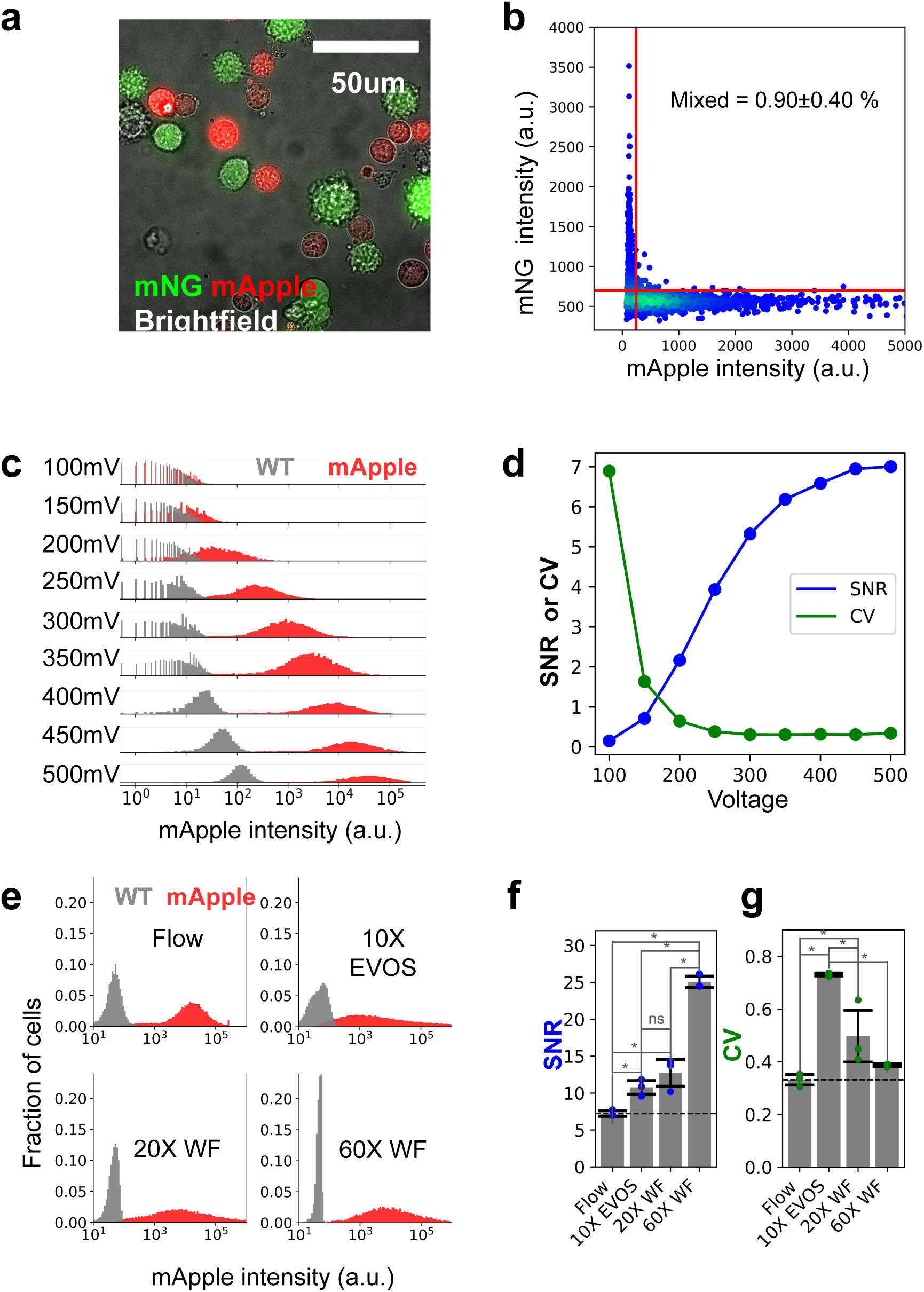

Next, we compared the specificity of our method to that of flow cytometry. We analyzed HEK cells stably expressing the fluorescent protein mApple and nonfluorescent wild-type (WT) HEK cells. To ensure we are making a fair comparison between flow cytometry and our imaging systems, we optimized the flow cytometer’s PMT voltage setting to maximize the signal-to-noise ratio (SNR) and minimize the coefficient of variation (CV) in mApple signal (Fig. 4c-d). We chose a PMT voltage of 450 mV as it had a good SNR, and increasing the voltage did not improve the CV (Fig. 4d). We compared SNR and CV values collected with these optimized flow settings to our approach utilizing two microscopes, including a more accessible, low-cost EVOS microscope. With the same cell population that we measured with flow cytometry, we captured images on our Nikon Ti-E widefield microscope with 20X and 60X magnification and on an EVOS microscope with 10X magnification. The distribution of mApple signal and WT background was comparable on all systems, with the notable exception that the width of the WT distribution for data obtained with the Nikon Ti-E was noticeably less than data obtained with flow or the EVOS microscope (Fig. 4e). We quantified this observation by calculating SNR for all four approaches. All imaging methods had an SNR significantly more than that of optimized flow cytometry (Fig. 4f). We also compared the spread or variation in signals obtained from the mApple HEK cells as we suspected that our imaging-based method might lead to higher levels of variation in fluorescence intensities. Indeed, we observed higher variance in the fluorescence intensities for the data obtained with the EVOS and 20X Nikon Ti-E than with flow cytometry but did not observe a significant difference in CV between 60X Nikon Ti-E data and flow cytometry data (Fig. 4g).

### Imaging-based cytometry for prime editing optimization

Prime editing is a genetic editing technique that utilizes a prime editor (PE), a single strand nicking Cas9 endonuclease fused to an engineered reverse transcriptase enzyme, a prime editing guide RNA (pegRNA) to edit or insert a new sequence into the genome(8), and a traditional single guide RNA (sgRNA) to facilitate opposing strand nicking. Instead of encoding base modification or insertions on exogenously added DNA, the genetic alterations in prime editing are encoded on the pegRNA and are copied into the genome via the reverse transcriptase (Fig. 5a). Prime editing does not make double-stranded breaks in the DNA like traditional CRISPR-Cas9 approaches, which are more error-prone(8). Along with the ability to edit post-mitotic cells and its potential to address almost 90% of disease-causing mutations, prime editing is an important technology in advancing genetic medicine. While prime editing is reasonably efficient at making small insertions up to 15 bases, the editing efficiency for inserts from 15 bases to 50 bases is inefficient and variable, depending on the genomic context(9). To improve prime editing, groups have optimized the molecular components of the prime editing system such as the pegRNA sequence(10) and the PE protein structure(11) and studied the endogenous DNA repair pathways involved in prime editing(12). It was shown that DNA mismatch repair (MMR) pathways strongly suppress the efficiency of prime editing. This information was used to develop a prime editing approach that inhibits MMR by transiently expressing a dominant negative MMR protein (MLH1dn)(12). Inefficiencies are also partially due to the suboptimal concentrations of the prime editing components (PE, pegRNA, sgRNA, MLH1dn). We reasoned that with the optimization of prime editing conditions, insertions of longer lengths could be made more reliable. One application that would benefit from efficient insertion is integration of the small strand of split fluorescent proteins for endogenous protein tagging. Endogenous protein tagging with split fluorescence proteins functions by genetically appending a small, ∼50 bp in most cases, tag to the coding region of a protein of interest. This results in the addition of amino acids of the small non-fluorescence fragment of a fluorescent protein being incorporated into the protein of interest. In the presence of the nonfluorescent large fragment, the small and large fragments combined to form the complete fluorescent form of a reporter(13).

**Fig. 5.**
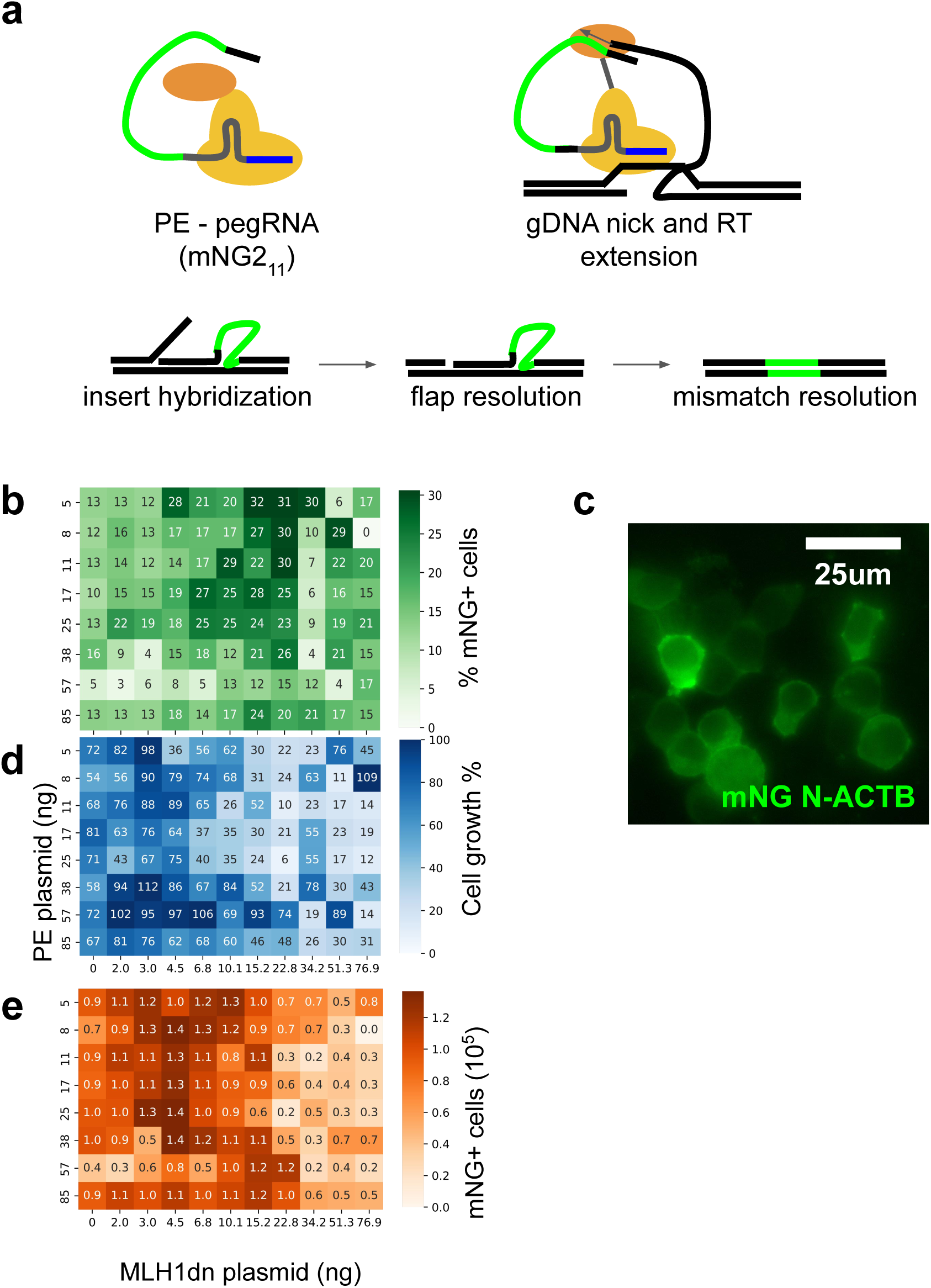
Prime editing optimization. **a**. Schematic of prime editing. PE and pegRNA complexes to form an active editor, the editor is targeted to specific location in genome via the pegRNA, the editor nicks the gDNA and pegRNA hybridizes to nicked strand, the reverse transcriptase on the editor extends the gDNA which includes the insert, the extended strand hybridizes to the opposite gDNA strand displacing the endogenous gDNA, the endogenous gDNA flap is excised, and the mismatched edit and endogenous gDNA is resolved. **b**. Heatmap of %mNG positive cells for all PE and MLH1dn conditions tested. **c**. Representative image of mNG tagged beta-actin **d**. Heatmap of relative cell proliferation for all PE and MLH1dn conditions tested. **e**. Heatmap of total mNG positive cells for all PE and MLH1dn conditions tested.

To demonstrate the utility of our microscopy cytometry method, we designed a pegRNA and sgRNA targeting the N-terminus of the beta-actin gene encoding an insertion of the mNG2_11_ fragment (48 bp). We screened a set of 88 prime editing conditions utilizing positive mNG fluorescence as a readout of prime editing insertion of mNG2_11_. We varied the PE plasmid concentrations and the MLH1dn plasmid concentration and assessed editing efficiency after five days. We calculated the percent of mNG positive cells and observed that a PE plasmid concentration of 5 ng and MLH1dn plasmid concentration of 15.2 ng produced the highest fraction of mNG positive cells (Fig. 5b). With the same MLH1dn condition, increasing the amount of loaded PE plasmid decreased the fraction of mNG-positive cells (Fig. 5b). We inspected our images for proper localization of mNG signal in cells and observed that it was predominantly in the cytoplasm (Fig. 5c), consistent with previous reports of mNG2_11_ tagging of beta-actin(14). We noticed fewer total cells in conditions with high concentrations of MLH1dn. To quantify this effect, we counted the number of Cellpose segments for each condition and normalized it to the no plasmid control. We observed that MLH1dn inhibited cell proliferation in a dose-dependent manner, with the lowest relative cell numbers observed for the highest concentration of MLH1dn (Fig. 5d). For endogenous gene tagging, the total number of mNG-positive cells is a practical metric to assess for cell editing throughput as it integrates editing efficiency and cell proliferation. We counted the total number of mNG positive cells for each condition and found that a lower MLH1dn concentration of 4.5 ng yielded the most mNG positive cells, with PE concentration having little effect on the total number of mNG positive cells (Fig. 5e).

## Discussion

Microscopy-based cytometry, as we demonstrated here, has multiple benefits. First, the use of imaging approaches provides richer information per cell than flow cytometers, which can be further analyzed after the fact, which is not the case with flow cytometry. Second, imaging approaches allow for minimal handling of cells, only requiring a brief exposure to trypsin as sample prep, which could be easily automated. Third, this approach only requires a simple microscope, making it accessible to many researchers.

Microscopy-based cytometry relies heavily on the ability to segment cells from the background and other cells accurately. While Cellpose may work well for many cell types and conditions, particularly for fluorescent images, in our hands, it failed to reliably segment HEK cells adhered to the surface using brightfield images only (Fig. 1a,d). This poor performance is most likely due to the data type used to train the segmentation model and low contrast from brightfield images of adherent cells. By inducing slight detachment of the cell from the surface with trypsin, the contrast at the edges increased, allowing Cellpose to demark the boundaries of cells more accurately (Fig. 1b,d). When cells were completely detached from the surface, Cellpose can over-segment the images, providing more segments than real cells (Fig. 1c-d). Cellpose performing better on only partially detached cells is consistent with the surface adhered cell Cellpose was trained on as these partially adhered cells more resemble the training data. The mechanical disruption and centrifugation of cells led to some cells being stacked on each other, which could lead to misidentification due to out-of-focus fluorescence contributing to adjacent cells (Fig. 3a 125K).

From a practical perspective, the preparation of trypsinized-only cells is more streamlined than trypsinized and resuspended (also the typical workflow for flow cytometry), only requiring the addition of trypsin and a quencher before imaging, decreasing the time and complexity of experiments. A trypsinization-only method will maintain a cell’s relative location to other cells. This could allow colonies of cells to be assessed, facilitating a more complex analysis of many cells from a clonal subpopulation, which could be informative to understanding heterogeneity in biological processes. Additionally, because cells don’t completely detach and ball up with a trypsinization-only approach, if a nuclear stain is included, the cytoplasmic signal can easily be differentiated from the nuclear signal, which is not possible when cells are trypsinized and resuspended.

Cellpose parameter optimization demonstrated that proper selection of estimated diameters is important for accurate discrimination of cell boundaries, whereas the flow threshold had much less of an effect for the values we tested in our data (Fig. 2a-i). Cellpose does contain functionality to estimate a diameter parameter instead of using a user-defined value but for our data, Cellpose greatly underestimated the diameter and location of cells when using this functionality. Cellpose estimated the average diameter of cells in our data to be 23.76 ± 3.25 pixels which is well below the 145 pixels we found to be optimal. We suggest users of this method always be conscious of selecting an appropriate diameter parameters. Our observation that Cellpose will over-segment images of trypsinized and resuspended cells if improper parameters are chosen again suggests that the partially adhered cells more resemble the training data as the maximum segments obtained from this group was the same as the number of nuclei counted (Fig. 2b-c). For these reasons, we opted for the slight under-segmentation of trypsinized cells instead of fully trypsinized and resuspended cells in our method. The false negative and false positive ratios were calculated using the nuclei segments as the ground truth. This is a limited method as the nucleus does not encompass all of the cytoplasm, particularly in the adhered cells, but instead occupies 80-90% in our HEK cell images. For our purpose, these metrics are sufficient to capture when multiple segments are called for the same cell and when no segments are called for a cell.

We demonstrated that Cellpose faithfully segments all of the cells imaged up to 62.5k cells/well (Fig. 3b), beyond which the cell density was too high, leading to cells stacking on top of one another and underdetection. This demonstrates a cell density limitation to this approach, as high density and cells that grow in more than a single layer might negatively affect the accuracy of measurements. While we demonstrated that Cellpose works well for partially adhered cells, it is also practical to use with non-adherent cells if the cells are allowed to settle or be centrifuged to the surface. To overcome the limitation of Cellpose discussed, it is possible to add annotations to a Cellpose model and improve its accuracy in a future version of Cellpose(15). This could enhance segmentation in the case of non-adhered cells, but we found the pre-trained models sufficient for our purposes.

We saw very few segments that contained a substantial amount of signal from both mApple and mNG in our mixing experiments, which indicates a high level of single-cell purity (Fig 4a-b). This is an important metric to assess an imaging approach especially when comparing to flow cytometry. We suspected that cells in close proximity to each other would contribute to the fluorescence measurement of neighboring cells but we do not see a substantial influence of this. Our experiments comparing flow cytometry to our approach demonstrate that imaging can yield higher SNR measurements than a conventional flow cytometry approach.

This is likely due to microscopes’ optics, which are optimized better for light collection, and the amount of exposure time in which signal can be collected. For flow cytometry, the exposure can be on the order of less than a millisecond, while the exposure for microscopy is typically hundreds of milliseconds. This longer exposure time increases the signal while the noise remains relatively constant. While the laser power of the flow cytometer could always be increased to compensate for shorter exposure, this may saturate the emitting ability of fluorescence elements and could induce phototoxicity(16). Imaging-based approaches, as we demonstrated, have the ability to measure lowly expressed proteins which can not be measured on most flow cytometers. This has been shown to be particularly useful in assessing endogenous protein tagging efficiency(17) and allows for a larger dynamic range in measurements of fluorescent reporters.

A typical flow cytometry experiment takes approximately 2 hours to process a 96-well plate with 50,000 cells per well, which includes 0.5 hours for the cell preparation and 1.5 hours of flow cytometer time. The throughput of microscopy-based cytometry largely depends on the size of the microscope field of view and the automation of the microscope. At a maximum cell density of 62.5k cells per well of a 96-well plate (∼200k cells/cm2), using 10X magnification on a typical sCMOS camera (∼ 1.3 x 1.3 mm2 field of view), one image will contain approximately 3,300 cells therefore requiring 16 images per well to obtain the same number of cells as a flow cytometer. A standard microscope with a motorized stage can easily image a full 96-well plate in hours, closely matching the throughput of a flow cytometer. Additionally, due to the high level of coincident when high flow rates are used in flow cytometry, many events are discarded due to doublet formation increasing the time and number of cells required to complete a flow cytometry experiment.

We demonstrated the utility of our approach by screening a set of 88 prime editing conditions. We found an optimal concentration for the amount of PE and MLH1dn to use when tagging beta-actin with mNG_11_. The conditions with the highest fraction of mNG-positive cells (Fig. 5b) were ones with high levels of MLH1dn plasmids as well as some of the lowest cell proliferation (Fig. 5d). This slow cell growth is not surprising as MLH1 is involved in meiotic crossing over during cell proliferation(18) and MMR processes. While MMR inhibition with MLH1dn may promote prime editing even at low concentrations of PE plasmid, its impact on cell viability needs to be considered in prime editing optimization. We evaluated a more inclusive metric that considered both editing efficiency and cell proliferation (Fig. 5e) as it is more relevant to endogenous gene tagging methods with the ultimate goal of sorting cells into a pure population of tagged cells. This is much more efficient with more tagged cells in total. Here, we screened PE and MLH1dn plasmid concentrations but this method could easily be used to screen other relevant factors in prime editing such as pegRNA sequence, pegRNA targeting site, and novel PE architectures.

Beyond prime editing, this approach has utility in any assay measuring a fluorescent reporter that one would typically be assayed on a flow cytometer. With a better SNR, simpler sample preparation compared to flow cytometry, and richer data, even on simple microscopes, we envision this approach replacing flow cytometry for diagnostic assays and in situations where flow cytometers are not accessible. Additionally, this method is particularly useful where many samples require assaying, and automated sampling for a flow cytometer is not available as sample preparation is easily parallelizable.

## Methods

### Cell culture

HEK293 cells were cultured in DMEM + 10% FBS & 1% Penicillin-Streptomycin. Cells were washed twice with 1x PBS, then detached by incubating 2-5 minutes at room temperature with 3 mL of 0.25% trypsin. Cells were seeded on glass bottom 96-well plates. For Cellpose validation experiments, cells were stained with 10 μg/ml of Hoechst in PBS for 10 minutes. To induce partial detachment of cells facilitating better segmentation, cells were first washed with PBS and then treated with 0.25% trypsin for 3 minutes at 37°C. trypsin was quenched with DMEM + 10% FBS and replaced with FluoroBrite DMEM + 10% FBS for imaging. For the trypsinized and resuspended condition, cells were treated with 0.25% trypsin for 3 minutes at 37°C, quenched with DMEM + 10% FBS, and mechanically agitated by repeatedly mixing via pipetting the solution in the wells. Cells were then centrifuged in the wells for 3 minutes at 300 g to place cells in the imaging plane. The media/trypsin solution was gently removed and replaced with FluoroBrite DMEM + 10% FBS before imaging. For cell dilution experiments, HEK cells in suspension were counted and diluted to appropriate concentrations such that 500000, 250000, 125000, 62500, 31250, or 15625 cells were placed in each well. Cells were then centrifuged in the wells for 3 minutes at 300 g to place cells in the imaging plane. The media/trypsin solution was gently removed and replaced with FluoroBrite DMEM + 10% FBS before imaging. For cell mixing experiments, cells were mixed at appropriate ratio before being seeded and allowed to appear and grow for two days.

### Image acquisition

Samples were imaged with a widefield Nikon Ti-E microscope with a motorized stage, a Hamamatsu ORCA Flash 4.0 camera, an LED light source (Excelitas X-Cite XLED1), and a 60X CFI Plan Apo IR water immersion objective or a plan fluor 20X/0.45 air objective.

For SNR and CV comparisons, samples were also imaged with an EVOS Cell Imaging System (Thermo Fisher: AMAFD1000), RFP (Thermo Fisher: AMEP4652) EVOS LED filter cube, and a Plan Fluor 10X/0.3 air objective.

### Cellpose segmentation

Brightfield images were segmented with Cellpose (4) using the general cytoplasmic model described in the original publication. Segments on the edge of the images were removed as well as segments above or below thresholds for cell size. These thresholds were set by first discarding very small segments, in our case segments less than 2000 pixels for 60x images, then fitting a normal distribution to the remaining segment sizes and setting the size thresholds as two standard deviations away from the estimated mean in both directions. A similar approach was applied to fluorescent nuclei images using the nuclei model in Cellpose and these segments were used as a reference for calculating the Cellpose segments/ nuclei count and the false positive and false negative rates. The segmentation of nuclei with Cellpose is much more robust as the nuclei model in Cellpose was trained entirely on fluorescence images of nuclei whereas the cytoplasmic model was trained on both fluorescence and brightfield images.

Moreover, nuclei of adjacent cells are often spatially separated in the fluorescence images.

### Calculating Cellpose accuracy and optimal parameters

To calculate the number of Cellpose segments/nuclei the ratio of the number of Cellpose segments obtained from brightfield images after size thresholding was compared to the number of segments obtained from nuclei images after size thresholding for each image in each condition. To calculate the false negative ratio for Cellpose in each image, the number of brightfield segments that fell within each nuclei segment were calculated. If there were no brightfield segments, this segment was considered to be false negative. Similarly, to calculate the false positive ratio, the number of nuclei segments that fell within each brightfield segment were calculated. If there were no nuclei segments, this segment was considered a false positive.

To optimize the parameters for Cellpose, four representative images were considered for the adhered, trypsinized, and trypsinized/resuspended conditions. Cellpose was applied to all 12 images for each combination of 25 diameters ranging from 25 to 505 pixels and 21 flow threshold values from 0.05 to 2.05 for a total of 525 parameter combinations. The segments/nuclei, false positive ratio, and false negative ratios were calculated for each condition and set of parameters.

### Fluorescence cell line generation

To generate the landing pad cell line, HEK293 cells were transfected with three plasmids using jetOPTIMUS according to the manufacturer’s instructions in a 6 well plate with cells grown to 80% confluency. The plasmids used are as followed: 500 ng of a plasmid expressing Sniper2L Cas9 (19), 500 ng of a plasmid expressing a single guide RNA targeting the safe harbor locus in the first intron of the adeno-associated virus integration site 1 (AAVS1) (targeting sequence GTTAATGTGGCTCTGGTTCT) (Supplementary materials), and 3000 ng of a donor plasmid containing in order: 800 bp section of homology to the endogenous AAVS1 site, a splice acceptor, a T2A site, an attB site of the large serine recombinase Pa01 (5), a TagBFP-NLS, a T2A site, a puromycin resistance gene, a polyadenylation signal, and another 800bp section of homology to the endogenous AAVS1 site (Supplementary materials). After 5 days of editing, edited cells were enriched by continual exposure to 1ug/ml puromycin.

To integrate the mApple sequence into the AAVS1 site of landing pad cells, two plasmids were transfected in a similar fashion as above; 3000 ng of a plasmid containing an attP site of the Pa01 protein with an mApple expression cassette, and 500 ng of a plasmid to express the large serine recombinase Pa01 to facilitate mApple integration (Addgene plasmid #193460).

Integration was allowed to proceed for 5 days before mApple+, TagBFP- cells were enriched using fluorescence-activated cell sorting (FACS). HSPA8 mNeonGreen HeLa cells were generated as described previously (14).

### Cell fluorescence analysis

To quantify fluorescence signals from individual cells, the brightfield segmentation boundaries from Cellpose were converted to a mask and applied to all channels to exclude all data outside of segments, then for each unique segment, the sum of all the pixel values in each channel were calculated and normalized to the total number of pixels. Cells were considered positive for fluorescence signals in each channel if the average pixel intensity was greater than the 99.9% percentile value calculated for the parent cell line with no fluorescence protein. SNR was calculated by dividing the difference in mean fluorescence of mApple-positive and non-fluorescent cells by the standard deviation of non-fluorescent cells. CV was calculated by dividing the standard deviation of the fluorescence of mApple-positive cells by the difference in mean fluorescence of mApple-positive and non-fluorescent cells.

### Flow Cytometry

Flow cytometry was performed on a BD FACSAria II. The mNG signal was measured with a 488 nm laser and a 530/30 bandpass filter. The mApple signal was measured using the 561 nm laser and a 610/20 bandpass filter. For detector voltage optimization, the photomultiplier tube voltage was set at varying values and data collected for each voltage setting. 10,000 events were recorded for each sample. Files in the .fcs format were exported from the BD FACS Aria II and were analyzed in Python.

### Prime editing optimization

HEK293 cells stably expressing the mNG2_1-10_ fragment were seeded into a 96-well plate as described above. All cells were transfected with 15 ng of plasmids expressing sgRNA (protospacer: GAAGCCGGCCTTGCACATGC) and 15 ng of a plasmid expressing pegRNA targeting the N terminus of the beta-actin gene carrying an insert of mNG2_11_ (Supplementary materials) For the combination of conditions, we screened 11 MLH1dn plasmid (Addgene plasmid #174824) concentrations from 0 ng/well to 77 ng/well and 8 PE plasmid (Supplementary materials) concentrations from 5ng/well to 85ng/well. JetOPTIMUS was used for all transfections according to the manufacturer’s instructions. Cells were trypsinized to partially detached cells as described above and images were taken on day 5. Editing efficiency was calculated by taking the ratio of mNG-positive cells to total cells. Cell survival ratios were obtained by counting Cellpose segments compared to no plasmid controls.

### Statistical analysis

When error bars are present, data represents >= 3 independent experiments. To determine significance in the differences between groups analysis of variance (ANOVA) was performed and if significant Tukey HSD test was performed to assess significance between groups. significance was determined if p < 0.05

## Acknowledgments

This work is supported by the National Institutes of Health (R01GM131641 to B.H.). B.H. is a Chan Zuckerberg Biohub San Francisco Investigator.

## Author contributions

D.F. performed the experiments, analyzed the data, and contributed to the interpretation of the results. Y.K. performed flow cytometry experiments. K.Y. conceived and designed the proof-of-concept experiments. S.R. assisted in developing data analysis methods. D. F and

B.H. conceived the project. B.H. and D.F. wrote the manuscript.

## Data availability

A public git repository is available at https://github.com/BoHuangLab/dcyto containing source code and a minimum analytical dataset sufficient to reproduce figures in the main text.

