## Supplementary Information for "Accessible and accurate cytometry analysis using fluorescence microscopes"

### Microscopy Based Cytometry: SUPPLEMENTARY MATERIALS

Plasmid sequences

**AASV1_Pa01attb_BFP_puro_HA**
ctagtcctgcaggtttaaacgaattcgccctttgctttctctgaccagcattctctcccctgggcctgtgccgctttctgtctgcagcttgtggc
ctgggtcacctctacggctggcccagatccttccctgccgcctccttcaggttccgtcttcctccactccctcttccccttgctctctgctgtgtt
gctgcccaaggatgctctttccggagcacttccttctcggcgctgcaccacgtgatgtcctctgagcggatcctccccgtgtctgggtcct
ctccgggcatctctcctccctcacccaaccccatgccgtcttcactcgctgggttcccttttccttctccttctggggcctgtgccatctctcgtt
tcttaggatggccttctccgacggatgtctcccttgcgtcccgcctccccttcttgtaggcctgcatcatcaccgtttttctggacaaccccaa
agtaccccgtctccctggctttagccacctctccatcctcttgctttctttgcctggacaccccgttctcctgtggattcgggtcacctctcact
cctttcatttgggcagctcccctacccccccttacctctctagtctgtgctagctcttccagccccctgtcatggcatcttccaggggtccgag
agctcagctagtcttcttcctccaacccgggcccctatgtccacttcaggacagcatgtttgctgcctccagggatcctgtgtccccgagc
tgggaccaccttatattcccagggccggttaatgtggctctggttctgggtacttttatctgtcccctccaccccacagtggggcaagcttct
gacctcttctcttcctcccacagggcctcgagagatctggcagcggagagggcagaggaagtcttctaacatgcggtgacgtggagg
agaatcccggccctaggctcgagGAGACCGTGACCTACATGCTCGAAGGGCGTATGCGCCACGAAA
TGAGCGAGCTGATTAAGGAGAACATGCACATGAAGCTGTACATGGAGGGCACCGTGGACA
ACCATCACTTCAAGTGCACATCCGAGGGCGAAGGCAAGCCCTACGAGGGCACCCAGACCA
TGAGAATCAAGGTGGTCGAGGGCGGCCCTCTCCCCTTCGCCTTCGACATCCTGGCTACTA
GCTTCCTCTACGGCAGCAAGACCTTCATCAACCACACCCAGGGCATCCCCGACTTCTTCAA
GCAGTCCTTCCCTGAGGGCTTCACATGGGAGAGAGTCACCACATACGAAGACGGGGGCG
TGCTGACCGCTACCCAGGACACCAGCCTCCAGGACGGCTGCCTCATCTACAACGTCAAGA
TCAGAGGGGTGAACTTCACATCCAACGGCCCTGTGATGCAGAAGAAAACACTCGGCTGGG
AGGCCTTCACCGAAACGCTGTACCCCGCTGACGGCGGCCTGGAAGGCAGAAACGACATG
GCCCTGAAGCTCGTGGGCGGGAGCCATCTGATCGCAAACATCAAGACCACATATAGATCC
AAGAAACCCGCTAAGAACCTCAAGATGCCTGGCGTCTACTATGTGGACTACAGACTGGAAA
GAATCAAGGAGGCCAACAACGAAACCTACGTCGAGCAGCACGAGGTGGCAGTGGCCAGA
TACTGCGACCTCCCTAGCAAACTGGGGCACAAGCTTAATCCCAAGAAGAAGAGGAAGGTG
gagggcagaggaagtcttctaacatgcggtgacgtggaggagaatcccggccctaggctcgagatgaccgagtacaagcccacg
gtgcgcctcgccacccgcgacgacgtccccagggccgtacgcaccctcgccgccgcgttcgccgactaccccgccacgcgccac
accgtcgatccggaccgccacatcgagcgggtcaccgagctgcaagaactcttcctcacgcgcgtcgggctcgacatcggcaaggt
gtgggtcgcggacgacggcgccgcggtggcggtctggaccacgccggagagcgtcgaagcgggggcggtgttcgccgagatcg
gcccgcgcatggccgagttgagcggttcccggctggccgcgcagcaacagatggaaggcctcctggcgccgcaccggcccaagg
agcccgcgtggttcctggccaccgtcggcgtctcgcccgaccaccagggcaagggtctgggcagcgccgtcgtgctccccggagtg
gaggcggccgagcgcgccggggtgcccgccttcctggagacctccgcgccccgcaacctccccttctacgagcggctcggcttcac
cgtcaccgccgacgtcgaggtgcccgaaggaccgcgcacctggtgcatgacccgcaagcccggtgcctgaCTCGAGCGAC
GCCTCGACTGTGCCTTCTAGTTGCCAGCCATCTGTTGTTTGCCCCTCCCCCGTGCCTTCCT
TGACCCTGGAAGGTGCCACTCCCACTGTCCTTTCCTAATAAAATGAGGAAATTGCATCGCA
TTGTCTGAGTAGGTGTCATTCTATTCTGGGGGGTGGGGTGGGGCAGGACAGCAAGGGGG
AGGATTGGGAAGACAATAGCAGGCATGCTGGGGATCCacgactgacaggattggtgacagaaaagcccc
atccttaggcctcctccttcctagtctcctgatattgggtctaacccccacctcctgttaggcagattccttatctggtgacacacccccatttc
ctggagccatctctctccttgccagaacctctaaggtttgcttacgatggagccagagaggatcctgggagggagagcttggcagggg
gtgggagggaagggggggatgcgtgacctgcccggttctcagtggccaccctgcgctaccctctcccagaacctgagctgctctgac
gcggctgtctggtgcgtttcactgatcctggtgctgcagcttccttacacttcccaagaggagaagcagtttggaaaaacaaaatcaga
ataagttggtcctgagttctaactttggctcttcacctttctagtccccaatttatattgttcctccgtgcgtcagttttacctgtgagataaggcc
agtagccagccccgtcctggcagggctgtggtgaggagggggtgtccgtgtggaaaactccctttgtgagaatggtgcgtcctaggt
gttcaccaggtcgtggccgcctctactccctttctctttctccatccttctttccttaaagagtccccagtgctatctgggacatattcctccgcc
cagagcagggtcccgcttccctaaggccctgctctgggcttctgggtttgagtccttggcaagcccaggagaggcgctcaggcttccct
gtccccccttcctcgtccaccatctcatgcccctggctctcctgccccttccctacaggggttcctggctctgctctaagggcgaattcgcgg
ccgctaaattcaattcgccctatagtgagtcgtattacaattcactggccgtcgttttacaacgtcgtgactgggaaaaccctggcgttacc
caacttaatcgccttgcagcacatccccctttcgccagctggcgtaatagcgaagaggcccgcaccgatcgcccttcccaacagttgc

gcagcctatacgtacggcagtttaaggtttacacctataaaagagagagccgttatcgtctgtttgtggatgtacagagtgatattattgac
acgccggggcgacggatggtgatccccctggccagtgcacgtctgctgtcagataaagtctcccgtgaactttacccggtggtgcatat
cggggatgaaagctggcgcatgatgaccaccgatatggccagtgtgccggtctccgttatcggggaagaagtggctgatctcagcca
ccgcgaaaatgacatcaaaaacgccattaacctgatgttctggggaatataaatgtcaggcatgagattatcaaaaaggatcttcacc
tagatccttttcacgtagaaagccagtccgcagaaacggtgctgaccccggatgaatgtcagctactgggctatctggacaaggaa
aacgcaagcgcaaagagaaagcaggtagcttgcagtgggcttacatggcgatagctagactgggcggttttatggacagcaagcg
aaccggaattgccagctggggcgccctctggtaaggttgggaagccctgcaaagtaaactggatggctttcttgccgccaaggatctg
atggcgcaggggatcaagctctgatcaagagacaggatgaggatcgtttcgcatgattgaacaagatggattgcacgcaggttctcc
ggccgcttgggtggagaggctattcggctatgactgggcacaacagacaatcggctgctctgatgccgccgtgttccggctgtcagcg
caggggcgcccggttctttttgtcaagaccgacctgtccggtgccctgaatgaactgcaagacgaggcagcgcggctatcgtggctgg
ccacgacgggcgttccttgcgcagctgtgctcgacgttgtcactgaagcgggaagggactggctgctattgggcgaagtgccggggc
aggatctcctgtcatctcaccttgctcctgccgagaaagtatccatcatggctgatgcaatgcggcggctgcatacgcttgatccggcta
cctgcccattcgaccaccaagcgaaacatcgcatcgagcgagcacgtactcggatggaagccggtcttgtcgatcaggatgatctgg
acgaagagcatcaggggctcgcgccagccgaactgttcgccaggctcaaggcgagcatgcccgacggcgaggatctcgtcgtga
cccatgcatggcgatgcctgcttgccgaatatcatggtggaaaatggccgcttttctggattcatcgactgtggccggctgggtgtggcg
gaccgctatcaggacatagcgttggctacccgtgatattgctgaagagcttggcggcgaatgggctgaccgcttcctcgtgctttacggt
atcgccgctcccgattcgcagcgcatcgccttctatcgccttcttgacgagttcttctgaattattaacgcttacaatttcctgatgcggtatttt
ctccttacgcatctgtgcggtatttcacaccgcatcaggtggcacttttcggggaaatgtgcgcggaacccctatttgtttatttttctaaata
cattcaaatatgtatccgctcatgagattatcaaaaaggatcttcacctagatccttttaaattaaaaatgaagttttaaatcaatctaaagt
atatatgagtaaacttggtctgacagttaccaatgcttaatcagtgaggcacctatctcagcgatctgtctatttcgttcatccatagttgcct
gactccccgtcgtgtagataactacgatacgggagggcttaccatctggccccagtgctgcaatgataccgcgagacccacgctcac
cggctccagatttatcagcaataaaccagccagccggaagggccgagcgcagaagtggtcctgcaactttatccgcctccatccagt
ctattaattgttgccgggaagctagagtaagtagttcgccagttaatagtttgcgcaacgttgttgccattgctacaggcatcgtggtgtca
cgctcgtcgtttggtatggcttcattcagctccggttcccaacgatcaaggcgagttacatgatcccccatgttgtgcaaaaaagcggtta
gctccttcggtcctccgatcgttgtcagaagtaagttggccgcagtgttatcactcatggttatggcagcactgcataattctcttactgtcat
gccatccgtaagatgcttttctgtgactggtgagtactcaaccaagtcattctgagaatagtgtatgcggcgaccgagttgctcttgcccg
gcgtcaatacgggataataccgcgccacatagcagaactttaaaagtgctcatcattggaaaacgttcttcggggcgaaaactctcaa
ggatcttaccgctgttgagatccagttcgatgtaacccactcgtgcacccaactgatcttcagcatcttttactttcaccagcgtttctgggtg
agcaaaaacaggaaggcaaaatgccgcaaaaaagggaataagggcgacacggaaatgttgaatactcatactcttcctttttcaat
attattgaagcatttatcagggttattgtctcatgaccaaaatcccttaacgtgagttttcgttccactgagcgtcagaccccgtagaaaag
atcaaaggatcttcttgagatcctttttttctgcgcgtaatctgctgcttgcaaacaaaaaaaccaccgctaccagcggtggtttgtttgccg
gatcaagagctaccaactctttttccgaaggtaactggcttcagcagagcgcagataccaaatactgttcttctagtgtagccgtagttag
gccaccacttcaagaactctgtagcaccgcctacatacctcgctctgctaatcctgttaccagtggctgctgccagtggcgataagtcgt
gtcttaccgggttggactcaagacgatagttaccggataaggcgcagcggtcgggctgaacggggggttcgtgcacacagcccagc
ttggagcgaacgacctacaccgaactgagatacctacagcgtgagctatgagaaagcgccacgcttcccgaagggagaaaggcg
gacaggtatccggtaagcggcagggtcggaacaggagagcgcacgagggagcttccagggggaaacgcctggtatctttatagtc
ctgtcgggtttcgccacctctgacttgagcgtcgatttttgtgatgctcgtcaggggggcggagcctatggaaaaacgccagcaacgcg
gcctttttacggttcctggccttttgctggccttttgctcacatgttctttcctgcgttatcccctgattctgtggataaccgtattaccgcctttgag
tgagctgataccgctcgccgcagccgaacgaccgagcgcagcgagtcagtgagcgaggaagcggaagagcgcccaatacgca
aaccgcctctccccgcgcgttggccgattcattaatgcagctggcacgacaggtttcccgactggaaagcgggcagtgagcgcaac
gcaattaatgtgagttagctcactcattaggcaccccaggctttacactttatgcttccggctcgtatgttgtgtggaattgtgagcggataa
caatttcacacaggaaacagctatgaccatgattacgccaagctcagaattaaccctcactaaaggga

**U6_AASV1_HA_sgRNA**
gaagatcctttgatcttttctacggggtctgacgctcagtggaacgaaaactcacgttaagggattttggtcatgagattatcaaaaagga
tcttcacctagatccttttaaattaaaaatgaagttttaaatcaatctaaagtatatatgagtaaacttggtctgacagttaccaatgcttaatc
agtgaggcacctatctcagcgatctgtctatttcgttcatccatagttgcctgactccccgtcgtgtagataactacgatacgggagggctt
accatctggccccagtgctgcaatgataccgcgagatccacgctcaccggctccagatttatcagcaataaaccagccagccggaa
gggccgagcgcagaagtggtcctgcaactttatccgcctccatccagtctattaattgttgccgggaagctagagtaagtagttcgcca
gttaatagtttgcgcaacgttgttgccattgctacaggcatcgtggtgtcacgctcgtcgtttggtatggcttcattcagctccggttcccaac
gatcaaggcgagttacatgatcccccatgttgtgcaaaaaagcggttagctccttcggtcctccgatcgttgtcagaagtaagttggccg

cagtgttatcactcatggttatggcagcactgcataattctcttactgtcatgccatccgtaagatgcttttctgtgactggtgagtactcaac
caagtcattctgagaatagtgtatgcggcgaccgagttgctcttgcccggcgtcaatacgggataataccgcgccacatagcagaact
ttaaaagtgctcatcattggaaaacgttcttcggggcgaaaactctcaaggatcttaccgctgttgagatccagttcgatgtaacccactc
gtgcacccaactgatcttcagcatcttttactttcaccagcgtttctgggtgagcaaaaacaggaaggcaaaatgccgcaaaaaaggg
aataagggcgacacggaaatgttgaatactcatactcttcctttttcaatattattgaagcatttatcagggttattgtctcatgagcggatac
atatttgaatgtatttagaaaaataaacaaataggggttccgcgcacatttccccgaaaagtgccacctgacgtcgctagctgtacaaa
aaagcaggctttaaaggaaccaattcagtcgactggatccggtaccaaggtcgggcaggaagagggcctatttcccatgattccttca
tatttgcatatacgatacaaggctgttagagagataattagaattaatttgactgtaaacacaaagatattagtacaaaatacgtgacgta
gaaagtaataatttcttgggtagtttgcagttttaaaattatgttttaaaatggactatcatatgcttaccgtaacttgaaagtatttcgatttcttg
gctttatatatcttgtggaaaggacgaaacaccgcaccccacagtggggccaCTgttttagagctagaaatagcaagttaaaataag
gctagtccgttatcaacttgaaaaagtggcaccgagtcggtgcttttttaagcttgggccgctcgaggtacctctctacatatgacatgtg
agcaaaaggccagcaaaaggccaggaaccgtaaaaaggccgcgttgctggcgtttttccataggctccgcccccctgacgagcat
cacaaaaatcgacgctcaagtcagaggtggcgaaacccgacaggactataaagataccaggcgtttccccctggaagctccctcgt
gcgctctcctgttccgaccctgccgcttaccggatacctgtccgcctttctcccttcgggaagcgtggcgctttctcatagctcacgctgtag
gtatctcagttcggtgtaggtcgttcgctccaagctgggctgtgtgcacgaaccccccgttcagcccgaccgctgcgccttatccggtaa
ctatcgtcttgagtccaacccggtaagacacgacttatcgccactggcagcagccactggtaacaggattagcagagcgaggtatgt
aggcggtgctacagagttcttgaagtggtggcctaactacggctacactagaagaacagtatttggtatctgcgctctgctgaagccag
ttaccttcggaaaaagagttggtagctcttgatccggcaaacaaaccaccgctggtagcggtggtttttttgtttgcaagcagcagattac
gcgcagaaaaaaaggatctcaa

**Pa01-attp.1-mApple-polyA**
AAAAGGACAATTACAAACAGGAATCGAATGCAACCGGCGCAGGAACACTGCCAGCGCATC
AACAATATTTTCACCTGAATCAGGATATTCTTCTAATACCTGGAATGCTGTTTTCCCGGGGA
TCGCAGTGGTGAGTAACCATGCATCATCAGGAGTACGGATAAAATGCTTGATGGTCGGAA
GAGGCATAAATTCCGTCAGCCAGTTTAGTCTGACCATCTCATCTGTAACATCATTGGCAAC
GCTACCTTTGCCATGTTTCAGAAACAACTCTGGCGCATCGGGCTTCCCATACAATCGATAG
ATTGTCGCACCTGATTGCCCGACATTATCGCGAGCCCATTTATACCCATATAAATCAGCATC
CATGTTGGAATTTAATCGCGGCCTGGAGCAAGACGTTTCCCGTTGAATATGGCTCATAACA
CCCCTTGTATTACTGTTTATGTAAGCAGACAGTTTTATTGTTCATGATGATATATTTTTATCTT
GTGCAATGTAACATCAGAGATTTTGAGACACAACGTGGCTTTGTTGAATAAATCGAACTTTT
GCTGAGTTGAAGGATCAGTCATGACCAAAATCCCTTAACGTGAGTTTTCGTTCCACTGAGC
GTCAGACCCCGTAGAAAAGATCAAAGGATCTTCTTGAGATCCTTTTTTTCTGCGCGTAATCT
GCTGCTTGCAAACAAAAAAACCACCGCTACCAGCGGTGGTTTGTTTGCCGGATCAAGAGCT
ACCAACTCTTTTTCCGAAGGTAACTGGCTTCAGCAGAGCGCAGATACCAAATACTGTTCTT
CTAGTGTAGCCGTAGTTAGGCCACCACTTCAAGAACTCTGTAGCACCGCCTACATACCTCG
CTCTGCTAATCCTGTTACCAGTGGCTGCTGCCAGTGGCGATAAGTCGTGTCTTACCGGGTT
GGACTCAAGACGATAGTTACCGGATAAGGCGCAGCGGTCGGGCTGAACGGGGGGTTCGT
GCACACAGCCCAGCTTGGAGCGAACGACCTACACCGAACTGAGATACCTACAGCGTGAGC
TATGAGAAAGCGCCACGCTTCCCGAAGGGAGAAAGGCGGACAGGTATCCGGTAAGCGGC
AGGGTCGGAACAGGAGAGCGCACGAGGGAGCTTCCAGGGGGAAACGCCTGGTATCTTTA
TAGTCCTGTCGGGTTTCGCCACCTCTGACTTGAGCGTCGATTTTTGTGATGCTCGTCAGGG
GGGCGGAGCCTATGGAAAAACGCCAGCAACGCGGCCTTTTTACGGTTCCTGGCCTTTTGC
TGGCCTTTTGCTCACATGTTCTTTCCTGCGTTATCCCCTGATTCTGTGGATAACCGTGCGG
CCGCCAATATAACTTCGTATAATGTATGCTATACGAAGTTATCCCTGAATTCGCATCTAGAC
TGAACTGGCCGATAATTGCAGACGAGGAGCATCGCCCTTCCCCGGCCCTCAGGTAAGAGG
ACCAAATACCGTAGCCGTTTCCAATTTCAGTCCTTTAGCGCCACCTGGTGCTAACTACTCTA
TCACGCTTTTATCCAATAACTACCTTTGTAAATGTAACGCTCTTCGAGAAAGCAGATTCTCA
TATCCATCTTGAGTCTTCTTTCTCGCAAGACAACACGAAATAGACACAGTCTCTTCCCTAGC
TGTACACTGtGCCGTGAGCAAGGGCGAGGAGAATAACATGGCCATCATCAAGGAGTTCATG
CGCTTCAAGGTGCACATGGAGGGCTCCGTGAACGGCCACGAGTTCGAGATCGAGGGCGA
GGGCGAGGGCCGCCCCTACGAGGCCTTTCAGACCGCTAAGCTGAAGGTGACCAAGGGTG

GCCCCCTGCCCTTCGCCTGGGACATCCTGTCCCCTCAGTTCATGTACGGCTCCAAGGTCT
ACATTAAGCACCCAGCCGACATCCCCGACTACTTCAAGCTGTCCTTCCCCGAGGGCTTCA
GGTGGGAGCGCGTGATGAACTTCGAGGACGGCGGCATTATTCACGTTAACCAGGACTCCT
CCCTGCAGGACGGCGTGTTCATCTACAAGGTGAAGCTGCGCGGCACCAACTTCCCCTCCG
ACGGCCCCGTAATGCAGAAGAAGACCATGGGCTGGGAGGCCTCCGAGGAGCGGATGTAC
CCCGAGGACGGCGCCCTGAAGAGCGAGATCAAGAAGAGGCTGAAGCTGAAGGACGGCG
GCCACTACGCCGCCGAGGTCAAGACCACCTACAAGGCCAAGAAGCCCGTGCAGCTGCCC
GGCGCCTACATCGTCGACATCAAGTTGGACATCGTGTCCCACAACGAGGACTACACCATC
GTGGAACAGTACGAACGCGCCGAGGGCCGCCACTCCACCGGCGGCATGGACGAGCTGTA
CAAGTAGCTCGAGCGACGCCTCGACTGTGCCTTCTAGTTGCCAGCCATCTGTTGTTTGCCC
CTCCCCCGTGCCTTCCTTGACCCTGGAAGGTGCCACTCCCACTGTCCTTTCCTAATAAAAT
GAGGAAATTGCATCGCATTGTCTGAGTAGGTGTCATTCTATTCTGGGGGGTGGGGTGGGG
CAGGACAGCAAGGGGGAGGATTGGGAAGACAATAGCAGGCATGCTGGGGATGCGGTGGG
CTCTATCCGAGCGGCCGCGTGTTACAACCAATTAACCAATTCTGATTAGAAAAACTCATCG
AGCATCAAATGAAACTGCAATTTATTCATATCAGGATTATCAATACCATATTTTTGAAAAAGC
CGTTTCTGTAATGAAGGAGAAAACTCACCGAGGCAGTTCCATAGGATGGCAAGATCCTGGT
ATCGGTCTGCGATTCCGACTCGTCCAACATCAATACAACCTATTAATTTCCCCTCGTCAAAA
ATAAGGTTATCAAGTGAGAAATCACCATGAGTGACGACTGAATCCGGTGAGAATGGCAAAA
GCTTATGCATTTCTTTCCAGACTTGTTCAACAGGCCAGCCATTACGCTCGTCATCAAAATCA
CTCGCATCAACCAAACCGTTATTCATTCGTGATTGCGCCTGAGCGAGGCGAAATACGCGAT
CGCTGTT

**NACTB_pegRNA**
gaagatcctttgatcttttctacggggtctgacgctcagtggaacgaaaactcacgttaagggattttggtcatgagattatcaaaaagga
tcttcacctagatccttttaaattaaaaatgaagttttaaatcaatctaaagtatatatgagtaaacttggtctgacagttaccaatgcttaatc
agtgaggcacctatctcagcgatctgtctatttcgttcatccatagttgcctgactccccgtcgtgtagataactacgatacgggagggctt
accatctggccccagtgctgcaatgataccgcgagatccacgctcaccggctccagatttatcagcaataaaccagccagccggaa
gggccgagcgcagaagtggtcctgcaactttatccgcctccatccagtctattaattgttgccgggaagctagagtaagtagttcgcca
gttaatagtttgcgcaacgttgttgccattgctacaggcatcgtggtgtcacgctcgtcgtttggtatggcttcattcagctccggttcccaac
gatcaaggcgagttacatgatcccccatgttgtgcaaaaaagcggttagctccttcggtcctccgatcgttgtcagaagtaagttggccg
cagtgttatcactcatggttatggcagcactgcataattctcttactgtcatgccatccgtaagatgcttttctgtgactggtgagtactcaac
caagtcattctgagaatagtgtatgcggcgaccgagttgctcttgcccggcgtcaatacgggataataccgcgccacatagcagaact
ttaaaagtgctcatcattggaaaacgttcttcggggcgaaaactctcaaggatcttaccgctgttgagatccagttcgatgtaacccactc
gtgcacccaactgatcttcagcatcttttactttcaccagcgtttctgggtgagcaaaaacaggaaggcaaaatgccgcaaaaaaggg
aataagggcgacacggaaatgttgaatactcatactcttcctttttcaatattattgaagcatttatcagggttattgtctcatgagcggatac
atatttgaatgtatttagaaaaataaacaaataggggttccgcgcacatttccccgaaaagtgccacctgacgtcgctagctgtacaaa
aaagcaggctttaaaggaaccaattcagtcgactggatccggtaccaaggtcgggcaggaagagggcctatttcccatgattccttca
tatttgcatatacgatacaaggctgttagagagataattagaattaatttgactgtaaacacaaagatattagtacaaaatacgtgacgta
gaaagtaataatttcttgggtagtttgcagttttaaaattatgttttaaaatggactatcatatgcttaccgtaacttgaaagtatttcgatttcttg
gctttatatatcttgtggaaaggacgaaacaccGCTATTCTCGCAGCTCACCAgttttagagctagaaatagcaagttaa
aataaggctagtccgttatcaacttgaaaaagtggcaccgagtcggtgcGACGAGCGCGGCGATATCATCATCCA
TGGtgccgccCATCATATCGGTAAAGGCCTTTTGCCACTCCTTGAAGTTGAGCTCGGTCATTG
AGCTGCGAGAAtttttttaagcttgggccgctcgaggtacctctctacatatgacatgtgagcaaaaggccagcaaaaggccag
gaaccgtaaaaaggccgcgttgctggcgtttttccataggctccgcccccctgacgagcatcacaaaaatcgacgctcaagtcagag
gtggcgaaacccgacaggactataaagataccaggcgtttccccctggaagctccctcgtgcgctctcctgttccgaccctgccgctta
ccggatacctgtccgcctttctcccttcgggaagcgtggcgctttctcatagctcacgctgtaggtatctcagttcggtgtaggtcgttcgct
ccaagctgggctgtgtgcacgaaccccccgttcagcccgaccgctgcgccttatccggtaactatcgtcttgagtccaacccggtaag
acacgacttatcgccactggcagcagccactggtaacaggattagcagagcgaggtatgtaggcggtgctacagagttcttgaagtg
gtggcctaactacggctacactagaagaacagtatttggtatctgcgctctgctgaagccagttaccttcggaaaaagagttggtagctc
ttgatccggcaaacaaaccaccgctggtagcggtggtttttttgtttgcaagcagcagattacgcgcagaaaaaaaggatctcaa

**U6_ACTB-N_sgRNA**
gaagatcctttgatcttttctacggggtctgacgctcagtggaacgaaaactcacgttaagggattttggtcatgagattatcaaaaagga
tcttcacctagatccttttaaattaaaaatgaagttttaaatcaatctaaagtatatatgagtaaacttggtctgacagttaccaatgcttaatc
agtgaggcacctatctcagcgatctgtctatttcgttcatccatagttgcctgactccccgtcgtgtagataactacgatacgggagggctt
accatctggccccagtgctgcaatgataccgcgagatccacgctcaccggctccagatttatcagcaataaaccagccagccggaa
gggccgagcgcagaagtggtcctgcaactttatccgcctccatccagtctattaattgttgccgggaagctagagtaagtagttcgcca
gttaatagtttgcgcaacgttgttgccattgctacaggcatcgtggtgtcacgctcgtcgtttggtatggcttcattcagctccggttcccaac
gatcaaggcgagttacatgatcccccatgttgtgcaaaaaagcggttagctccttcggtcctccgatcgttgtcagaagtaagttggccg
cagtgttatcactcatggttatggcagcactgcataattctcttactgtcatgccatccgtaagatgcttttctgtgactggtgagtactcaac
caagtcattctgagaatagtgtatgcggcgaccgagttgctcttgcccggcgtcaatacgggataataccgcgccacatagcagaact
ttaaaagtgctcatcattggaaaacgttcttcggggcgaaaactctcaaggatcttaccgctgttgagatccagttcgatgtaacccactc
gtgcacccaactgatcttcagcatcttttactttcaccagcgtttctgggtgagcaaaaacaggaaggcaaaatgccgcaaaaaaggg
aataagggcgacacggaaatgttgaatactcatactcttcctttttcaatattattgaagcatttatcagggttattgtctcatgagcggatac
atatttgaatgtatttagaaaaataaacaaataggggttccgcgcacatttccccgaaaagtgccacctgacgtcgctagctgtacaaa
aaagcaggctttaaaggaaccaattcagtcgactggatccggtaccaaggtcggcaggaagagggcctatttcccatgattccttca
tatttgcatatacgatacaaggctgttagagagataattagaattaatttgactgtaaacacaaagatattagtacaaaatacgtgacgta
gaaagtaataatttcttgggtagtttgcagttttaaaattatgttttaaaatggactatcatatgcttaccgtaacttgaaagtatttcgatttcttg
gctttatatatcttgtggaaaggacgaaacaccGCTATTCTCGCAGCTCACCAgttttagagctagaaatagcaagttaa
aataaggctagtccgttatcaacttgaaaaagtggcaccgagtcggtgcttttttaagcttgggccgctcgaggtacctctctacatatga
catgtgagcaaaaggccagcaaaaggccaggaaccgtaaaaaggccgcgttgctggcgtttttccataggctccgcccccctgacg
agcatcacaaaaatcgacgctcaagtcagaggtggcgaaacccgacaggactataaagataccaggcgtttccccctggaagctc
cctcgtgcgctctcctgttccgaccctgccgcttaccggatacctgtccgcctttctcccttcgggaagcgtggcgctttctcatagctcacg
ctgtaggtatctcagttcggtgtaggtcgttcgctccaagctgggctgtgtgcacgaaccccccgttcagcccgaccgctgcgccttatcc
ggtaactatcgtcttgagtccaacccggtaagacacgacttatcgccactggcagcagccactggtaacaggattagcagagcgag
gtatgtaggcggtgctacagagttcttgaagtggtggcctaactacggctacactagaagaacagtatttggtatctgcgctctgctgaa
gccagttaccttcggaaaaagagttggtagctcttgatccggcaaacaaaccaccgctggtagcggtggtttttttgtttgcaagcagca
gattacgcgcagaaaaaaaggatctcaa

**pCMV-PEmax-P2A**
gacattgattattgactagttattaatagtaatcaattacggggtcattagttcatagcccatatatggagttccgcgttacataacttacggt
aaatggcccgcctggctgaccgcccaacgacccccgcccattgacgtcaataatgacgtatgttcccatagtaacgccaatagggac
tttccattgacgtcaatgggtggagtatttacggtaaactgcccacttggcagtacatcaagtgtatcatatgccaagtacgccccctattg
acgtcaatgacggtaaatggcccgcctggcattatgcccagtacatgaccttatgggactttcctacttggcagtacatctacgtattagt
catcgctattaccatggtgatgcggttttggcagtacatcaatgggcgtggatagcggtttgactcacggggatttccaagtctccacccc
attgacgtcaatgggagtttgttttggcaccaaaatcaacgggactttccaaaatgtcgtaacaactccgccccattgacgcaaatggg
cggtaggcgtgtacggtgggaggtctatataagcagagctggtttagtgaaccgtcagatccgctagagatccgcggccgctaatac
gactcactatagggagagccgccaccatgaaacggacagccgacggaagcgagttcgagtcaccaaagaagaagcggaaagt
cgacaagaagtacagcatcggcctggacatcggcaccaactctgtgggctgggccgtgatcaccgacgagtacaaggtgcccagc
aagaaattcaaggtgctgggcaacaccgaccggcacagcatcaagaagaacctgatcggagccctgctgttcgacagcggcgaa
acagccgaggccacccggctgaagagaaccgccagaagaagatacaccagacggaagaaccggatctgctatctgcaagaga
tcttcagcaacgagatggccaaggtggacgacagcttcttccacagactggaagagtccttcctggtggaagaggataagaagcac
gagcggcaccccatcttcggcaacatcgtggacgaggtggcctaccacgagaagtaccccaccatctaccacctgagaaagaaa
ctggtggacagcaccgacaaggccgacctgcggctgatctatctggccctggcccacatgatcaagttccggggccacttcctgatcg
agggcgacctgaaccccgacaacagcgacgtggacaagctgttcatccagctggtgcagacctacaaccagctgttcgaggaaaa
ccccatcaacgccagcggcgtggacgccaaggccatcctgtctgccagactgagcaagagcagaaagctggaaaatctgatcgc
ccagctgcccggcgagaagaagaatggcctgttcggaaacctgattgccctgagcctgggcctgacccccaacttcaagagcaact
tcgacctggccgaggatgccaaactgcagctgagcaaggacacctacgacgacgacctggacaacctgctggcccagatcggcg
accagtacgccgacctgtttctggccgccaagaacctgtccgacgccatcctgctgagcgacatcctgagagtgaacaccgagatca

ccaaggccccctgagcgcctctatgatcaagagatacgacgagcaccaccaggacctgaccctgctgaaagctctcgtgcggca
gcagctgcctgagaagtacaaagagattttcttcgaccagagcaagaacggctacgccggctacattgacggcggagccagccag
gaagagttctacaagttcatcaagcccatcctggaaaagatggacggcaccgaggaactgctcgtgaagctgaagagagaggac
ctgctgcggaagcagcggaccttcgacaacggcagcatcccccaccagatccacctgggagagctgcacgccattctgcggcggc
aggaagatttttacccattcctgaaggacaaccgggaaaagatcgagaagatcctgaccttccgcatcccctactacgtgggccctct
ggccaggggaaacagcagattcgcctggatgaccagaaagagcgaggaaaccatcacccccctggaacttcgaggaagtggtgg
acaagggcgcttccgcccagagcttcatcgagcggatgaccaacttcgataagaacctgcccaacgagaaggtgctgcccaagca
cagcctgctgtacgagtacttcaccgtgtataacgagctgaccaaagtgaaatacgtgaccgagggaatgagaaagcccgccttcct
gagcggcgagcagaaaaaggccatcgtggacctgctgttcaagaccaaccggaaagtgaccgtgaagcagctgaaagaggact
acttcaagaaaatcgagtgcttcgactccgtggaaatctccggcgtggaagatcggttcaacgcctccctgggcacataccacgatct
gctgaaaattatcaaggacaaggacttcctggacaatgaggaaaacgaggacattctggaagatatcgtgctgaccctgacactgttt
gaggacagagagatgatcgaggaacggctgaaaacctatgcccacctgttcgacgacaaagtgatgaagcagctgaagcggcg
gagatacaccggctggggcaggctgagccggaagctgatcaacggcatccgggacaagcagtccggcaagacaatcctggattt
cctgaagtccgacggcttcgccaacagaaacttcatgcagctgatccacgacgacagcctgacctttaaagaggacatccagaaag
cccaggtgtccggccagggcgatagcctgcacgagcacattgccaatctggccggcagccccgccattaagaagggcatcctgca
gacagtgaaggtggtggacgagctcgtgaaagtgatgggccggcacaagcccgagaacatcgtgatcgaaatggccagagaga
accagaccacccagaagggacagaagaacagccgcgagagaatgaagcggatcgaagagggcatcaaagagctgggcagc
cagatcctgaaagaacaccccgtggaaaacacccagctgcagaacgagaagctgtacctgtactacctgcagaatgggcgggat
atgtacgtggaccaggaactggacatcaaccggctgtccgactacgatgtggacgctatcgtgcctcagagctttctgaaggacgact
ccatcgacaacaaggtgctgaccagaagcgacaagaaccggggcaagagcgacaacgtgccctccgaagaggtcgtgaagaa
gatgaagaactactggcggcagctgctgaacgccaagctgattacccagagaaagttcgacaatctgaccaaggccgagagagg
cggcctgagcgaactggataaggccggcttcatcaagagacagctggtggaaacccggcagatcacaaagcacgtggcacagat
cctggactcccggatgaacactaagtacgacgagaatgacaagctgatccgggaagtgaaagtgatcaccctgaagtccaagctg
gtgtccgatttccggaaggatttccagttttacaaagtgcgcgagatcaacaactaccaccacgcccacgacgcctacctgaacgcc
gtcgtgggaaccgccctgatcaaaaagtaccctaagctggaaagcgagttcgtgtacggcgactacaaggtgtacgacgtgcggaa
gatgatcgccaagagcgagcaggaaatcggcaaggctaccgccaagtacttcttctacagcaacatcatgaactttttcaagaccga
gattaccctggccaacggcgagatccggaagcggcctctgatcgagacaaacggcgaaaccggggagatcgtgtgggataaggg
ccgggattttgccaccgtgcggaaagtgctgagcatgccccaagtgaatatcgtgaaaaagaccgaggtgcagacaggcggcttca
gcaaagagtctatcctgcccaagaggaacagcgataagctgatcgccagaaagaaggactgggaccctaagaagtacggcggct
tcgacagccccaccgtggcctattctgtgctggtggtggccaaagtggaaaagggcaagtccaagaaactgaagagtgtgaaaga
gctgctggggatcaccatcatggaaagaagcagcttcgagaagaatcccatcgactttctggaagccaagggctacaaagaagtg
aaaaaggacctgatcatcaagctgcctaagtactccctgttcgagctggaaaacggccggaagagaatgctggcctctgccggcga
actgcagaagggaaacgaactggccctgccctccaaatatgtgaacttcctgtacctggccagccactatgagaagctgaagggctc
ccccgaggataatgagcagaaacagctgtttgtggaacagcacaagcactacctggacgagatcatcgagcagatcagcgagttct
ccaagagagtgatcctggccgacgctaatctggacaaagtgctgtccgcctacaacaagcaccgggataagcccatcagagagca
ggccgagaatatcatccacctgtttaccctgaccaatctgggagcccctgccgccttcaagtactttgacaccaccatcgaccggaag
aggtacaccagcaccaaagaggtgctggacgccaccctgatccaccagagcatcaccggcctgtacgagacacggatcgacctg
tctcagctgggaggtgactccggcggaagctctggtggcagcaagcggaccgccgacggctctgaattcgagagccctaagaaga
aaagaaaggtgagcggaggctctagcggcggaagcaccctgaacattgaagacgagtatagactgcatgaaacaagcaaggaa
cccgacgtgtccctgggctccacctggctgtccgactttccccaggcctgggccgagacaggaggaatgggcctggccgtgcggca
ggcaccccctgatcatccctctgaaggccacctctacacccgtgagcatcaagcagtaccctatgtctcaggaggccagactgggcat
caagcctcacatccagaggctgctggaccagggcatcctggtgccatgccagagcccctggaacacaccactgctgcccgtgaag
aagccaggcaccaatgactatagacccgtgcaggatctgagagaggtgaacaagagggtggaggatatccaccccaccgtgccc
aacccttacaatctgctgtccggcctgccccccttctcaccagtggtatacagtgctggacctgaaggatgccttcttttgtctgagactgca
ccctaccagccagccactgttcgcctttgagtggagggaccctgagatgggcatctctggccagctgacctggacacgcctgcctcag
ggcttcaagaatagcccaacactgtttaacgaggccctgcaccgcgacctggcagatttccggatccagcacccagatctgatcctgc
tgcagtacgtggacgatctgctgctggccgccaccagcgagctggattgccagcagggaacacgcgccctgctgcagaccctggg
aaacctgggatatagggcatccgccaagaaggcccagatctgtcagaagcaggtgaagtacctgggctatctgctgaaggagggc
cagagatggctgacagaggccaggaaggagacagtgatgggccagccaacacccaagaccccaagacagctgagggagttcc
tgggcaaagcaggattttgcaggctgttcatcccaggattcgcagagatggcagcacctctgtacccactgaccaagccgggcaccc
tgtttaattggggccctgaccagcagaaggcctatcaggagatcaagcaggccctgctgacagcaccagccctgggcctgccagac

ctgaccaagcctttcgagctgtttgtggatgagaagcagggctacgccaagggcgtgctgacccagaagctgggaccatggagacg
gcccgtggcctatctgtccaagaagctggacccagtggcagcaggatggccaccatgcctgaggatggtggcagcaatcgccgtgc
tgacaaaggatgccggcaagctgaccatgggacagccactggtcatcctggcaccacacgcagtggaggccctggtgaagcagc
ctccagatcgctggctgtctaacgcccggatgacacactaccaggccctgctgctggacaccgatcgcgtgcagtttggccctgtggtg
gccctgaatccagccaccctgctgcctctgccagaggagggcctgcagcacaactgtctggacatcctggcagaggcacacggaa
caaggccagacctgaccgatcagcccctgcctgacgccgatcacacatggtataccgatggaagctccctgctgcaggagggcca
gaggaaggcaggagcagcagtgaccacagagacagaagtgatctgggccaaggccctgccagcaggcacatccgcccagcg
ggccgagctgatcgccctgacccaggccctgaagatggccgagggcaagaagctgaacgtgtacacagactccagatatgccttc
gccaccgcacacatccacggagagatctacaggcgccggggctggctgacctctgagggcaaggagatcaagaacaaggatga
gatcctggccctgctgaaggccctgtttctgcccaagcggctgagcatcatccactgtcctggacaccagaagggacactccgccga
ggcaaggggcaatcggatggccgaccaggccgccagaaaggctgctattactgaaactcccgacacttccactctgctgattgaaa
actcctccccttctggcggctcaaaaagaaccgccgacggcagcgaattcgagtctcccaagaagaagaggaaagtcggctctgg
ccctgccgctaagagagtgaagctggacggatccggcgcaacaaacttctctctgctgaaacaagccggagatgtcgaagagaat
cctggaccgccgcatcatcaccatcaccattgagtttaaacccgctgatcagcctcgactgtgccttctagttgccagccatctgttgtttg
cccctcccccgtgccttccttgaccctggaaggtgccactcccactgtcctttcctaataaaatgagaaattgcatcgcattgtctgagta
ggtgtcattctattctggggggtggggtggggcaggacagcaagggggaggattgggaagacaatagcaggcatgctggggatgc
ggtgggctctatggcttctgaggcggaaagaaccagctggggctcgataccgtcgacctctagctagagcttggcgtaatcatggtcat
agctgtttcctgtgtgaaattgttatccgctcacaattccacacaacatacgagccggaagcataaagtgtaaagcctagggtgcctaat
gagtgagctaactcacattaattgcgttgcgctcactgcccgctttccagtcgggaaacctgtcgtgccagctgcattaatgaatcggcc
aacgcgcggggagaggcggtttgcgtattgggcgctcttccgcttcctcgctcactgactcgctgcgctcggtcgttcggctgcggcga
gcggtatcagctcactcaaaggcggtaatacggttatccacagaatcaggggataacgcaggaaagaacatgtgagcaaaaggc
cagcaaaaggccaggaaccgtaaaaaggccgcgttgctggcgtttttccataggctccgcccccctgacgagcatcacaaaaatcg
acgctcaagtcagaggtggcgaaacccgacaggactataaagataccaggcgtttccccctggaagctccctcgtgcgctctcctgtt
ccgaccctgccgcttaccggatacctgtccgcctttctcccttcgggaagcgtggcgctttctcatagctcacgctgtaggtatctcagttc
ggtgtaggtcgttcgctccaagctgggctgtgtgcacgaaccccccgttcagcccgaccgctgcgccttatccggtaactatcgtcttga
gtccaacccggtaagacacgacttatcgccactggcagcagccactggtaacaggattagcagagcgaggtatgtaggcggtgcta
cagagttcttgaagtggtggcctaactacggctacactagaagaacagtatttggtatctgcgctctgctgaagccagttaccttcggaa
aaagagttggtagctcttgatccggcaaacaaaccaccgctggtagcggtggtttttttgtttgcaagcagcagattacgcgcagaaaa
aaaggatctcaagaagatcctttgatcttttctacggggtctgacactcagtggaacgaaaactcacgttaagggattttggtcatgagat
tatcaaaaaggatcttcacctagatccttttaaattaaaaatgaagttttaaatcaatctaaagtatatatgagtaaacttggtctgacagtt
accaatgcttaatcagtgaggcacctatctcagcgatctgtctatttcgttcatccatagttgcctgactccccgtcgtgtagataactacga
tacgggagggcttaccatctggccccagtgctgcaatgataccgcgagacccacgctcaccggctccagatttatcagcaataaacc
agccagccggaagggccgagcgcagaagtggtcctgcaactttatccgcctccatccagtctattaattgttgccgggaagctagagt
aagtagttcgccagttaatagtttgcgcaacgttgttgccattgctacaggcatcgtggtgtcacgctcgtcgtttggtatggcttcattcagc
tccggttcccaacgatcaaggcgagttacatgatcccccatgttgtgcaaaaaagcggttagctccttcggtcctccgatcgttgtcaga
agtaagttggccgcagtgttatcactcatggttatggcagcactgcataattctcttactgtcatgccatccgtaagatgcttttctgtgactg
gtgagtactcaaccaagtcattctgagaatagtgtatgcggcgaccgagttgctcttgcccggcgtcaatacgggataataccgcgcc
acatagcagaactttaaaagtgctcatcattggaaaacgttcttcggggcgaaaactctcaaggatcttaccgctgttgagatccagttc
gatgtaacccactcgtgcacccaactgatcttcagcatcttttactttcaccagcgtttctgggtgagcaaaaacaggaaggcaaaatg
ccgcaaaaaagggaataagggcgacacggaaatgttgaatactcatactcttcctttttcaatattattgaagcatttatcagggttattgt
ctcatgagcggatacatatttgaatgtatttagaaaaataaacaaataggggttccgcgcacatttccccgaaaagtgccacctgacgt
cgacggatcgggagatcgatctcccgatcccctagggtcgactctcagtacaatctgctctgatgccgcatagttaagccagtatctgct
ccctgcttgtgtgttggaggtcgctgagtagtgcgcgagcaaaatttaagctacaacaaggcaaggcttgaccgacaattgcatgaag
aatctgcttagggttaggcgttttgcgctgcttcgcgatgtacgggccagatatacgcgtt
